# Development of noninvasive imaging to measure spontaneous pain in mice

**DOI:** 10.1101/2025.05.20.655170

**Authors:** Subo Yuan, Jia Yi Liew, Ajay Pal, Jigong Wang, Dustin P. Green, Massoud Motamedi, Shao-Jun Tang

## Abstract

Objectively measuring pain in laboratory animals is essential for pain research and analgesic development. Despite the development of various behavioral tests to measure evoked pain in animal models, measuring spontaneous pain remains challenging. To address this unmet need, we developed a novel imaging approach to detect spontaneous nociception in animal pain models. To do this, we generated a Bacterial Artificial Chromosomes transgenic mouse that expresses Redquorin under the murine synapsin 1 promoter. Redquorin is a fusion protein consisting of chimera and a 2x tandem dimer Tomato Aequorin (tdTA), which emits long wavelength bioluminescence from activated neurons in the presence of coelenterazine. This luminescence can penetrate tissues and form a projected image on the body surface that can be detected with a spectrum In Vivo Imaging System, thus creating a Nociceptive Neuronal Activity Imaging mouse. We used the tdTA mice to image bioluminescence in the spinal regions as a surrogate of spontaneous pain induced by capsaicin, the HIV-1 envelope glycoprotein gp120, and spinal nerve ligation. Results show that Redquorin-emitted bioluminescence is a sensitive optical surrogate to measure spontaneous pain. This approach offers a new method to measure spontaneous pain in animal models for basic and translational research.

## Introduction

Chronic pain is a highly subjective and unpleasant perception, characterized by a complex neuropathological pathogenesis within the somatosensory nervous system^[13; 32; 45]^. Chronic pain affects 21% of adults in the US according to the Centers for Disease Control and Prevention (2021), with 7% experiencing high-impact chronic pain^[34]^. However, only 30% of patients receive proper treatment^[9; 10]^. Despite increasing investment in pain research, the availability of new analgesics with ideal efficacy remains limited [7]. Many drug candidates show effective analgesia in animal models but then fail in clinical trials. One reason for this is likely due to experimental pain measurements focusing mainly on evoked pain rather than spontaneous pain, a natural component of chronic pain in patients. This misalignment between animal studies and human pain underscores the need for novel methodologies to measure spontaneous pain in animal models.

Animal pain models are developed to replicate clinical pain phenotypes, explore pain mechanisms, and evaluate the efficacy of potential drug candidates. Although evoked pain is commonly measured in animal models using reflexive behavior tests in response to mechanical (e.g., von Frey test) or temperature stimuli, there are limited also time consuming methods to evaluate spontaneous pain^[13; 20; 30],^. Measuring evoked pain is susceptible to experimenter bias and can be confounded by human-animal interactions^[30; 35]^. Animals can also memorize the pain-evoking stimuli and may thus exhibit avoidance behaviors, further complicating test results. Measuring pain with minimum experimenter handling may substantially reduce these confounders.Therefore, a platform to measure the effect of analgesics on spontaneous pain is highly desirable for pain basic research.

The activation of pain processing neurons elevates intracellular Ca^2+^, providing an opportunity to use calcium signals as a biomarker for nociception^[1; 23; 37]^. Intracellular Ca^2+^ elevation can be monitored using aequorin, a calcium-activated photoprotein that emits bioluminescent photons via the mechanism of bioluminescence resonance energy transfer (BRET)^[3; 6; 31; 49]^. In fact, aequorin-based methods have been used to monitor various physiological processes^[24; 31; 37]^. In this study, we sought to develop an aequorin-based strategy to monitor spontaneous pain in animal models. We generated a transgenic mouse that expresses a transgene encoding a fusion protein named Redquorin^[2]^, which consisted of chimera 2x tandem dimer Tomato Aequorin (tdTA) under the control of a murine synapsin 1 promoter (mSyn1)^[12; 29; 40]^. Increases in intracellular neuronal Ca^2+^ due to painful stimuli can lead to aequorin activity and bioluminescence emitting longwave red light from tdTA via BRET^[2; 16]^. The longwave light, which can penetrate tissues, can be captured by the Spectrum In Vivo Imaging System (IVIS), enabling visualization of nociception signals in the spinal dorsal horn (SDH), dorsal root ganglion (DRG), and nerves. Thus, these tdTA bearing mice serve as a new tool of Nociceptive Neuronal Activity Imaging (NNAI).

We generated three different pain models in the tdTA mice and measured photon emission from the spinal regions, using the IVIS imager. Our data provides proof-of-concept of this strategy and the application of this approach in measuring spontaneous pain in rodent models.

## Methods

### murine synapsin 1 promoter (mSyn1) Redquorin (tdTA) transgenic mouse

We constructed a fusion protein containing photoprotein tdTA and put it under mSyn1 to restrict its expression in neurons. Aequorin is a calcium-activated photoprotein which originally emits blue light when binding to Ca^2+^. A modified aequorin, the tdTA, produces a 575 nm emission shift through BRET and the emitted red light can cross the skin and other tissues with reduced photon scatter^[2; 4]^.

The transgenic mice were developed by bacterial artificial chromosomes (BAC) technique^[17]^. The Redquorin chimeric photoprotein was inserted under the mSyn1 promoter and the sequence was obtained from the gene bank, NM_001110780.1 and synapsin1 wide type allele located at chromosomal localization: X. <u>BAC Vector generation</u>: the Redquorin, short FRT sites flexed G418 (Geneticin) neo, and antibiotic selecting marker were inserted into exon 1 of synapsin 1 to generate a modification cassette (**Fig. 1A**). After site-directed recombination of flippase**-**flippase recognition, this target recombination mediated a removal of the Neo selection marker to construct the mSyn1-Redquorin-SV40pA cassette. The 5’ and 3’ homology arms, FRT site, Redquorin, and SV40pA in RP23-366P16-Redquorin-SV40pA were verified by DNA sequencing. The BAC transgenic mouse is C57BL6 × C57BL6. Among the 62 available pups, 5 were screened positive by the primers F1, CCTGCGTAAGCGGGGCACATTT and F2, GCCACTCTTTGGATGTTGGG, with a PCR product of 434bp. The RP23-366P16-Redquorin-SV40pA mouse was designed so that Redquorin chimeric protein expression was restricted to neurons. The mSyn1 Redquorin mouse was made under contract with Cyagen Biosciences (Santa Clara, CA).

**Fig 1.**
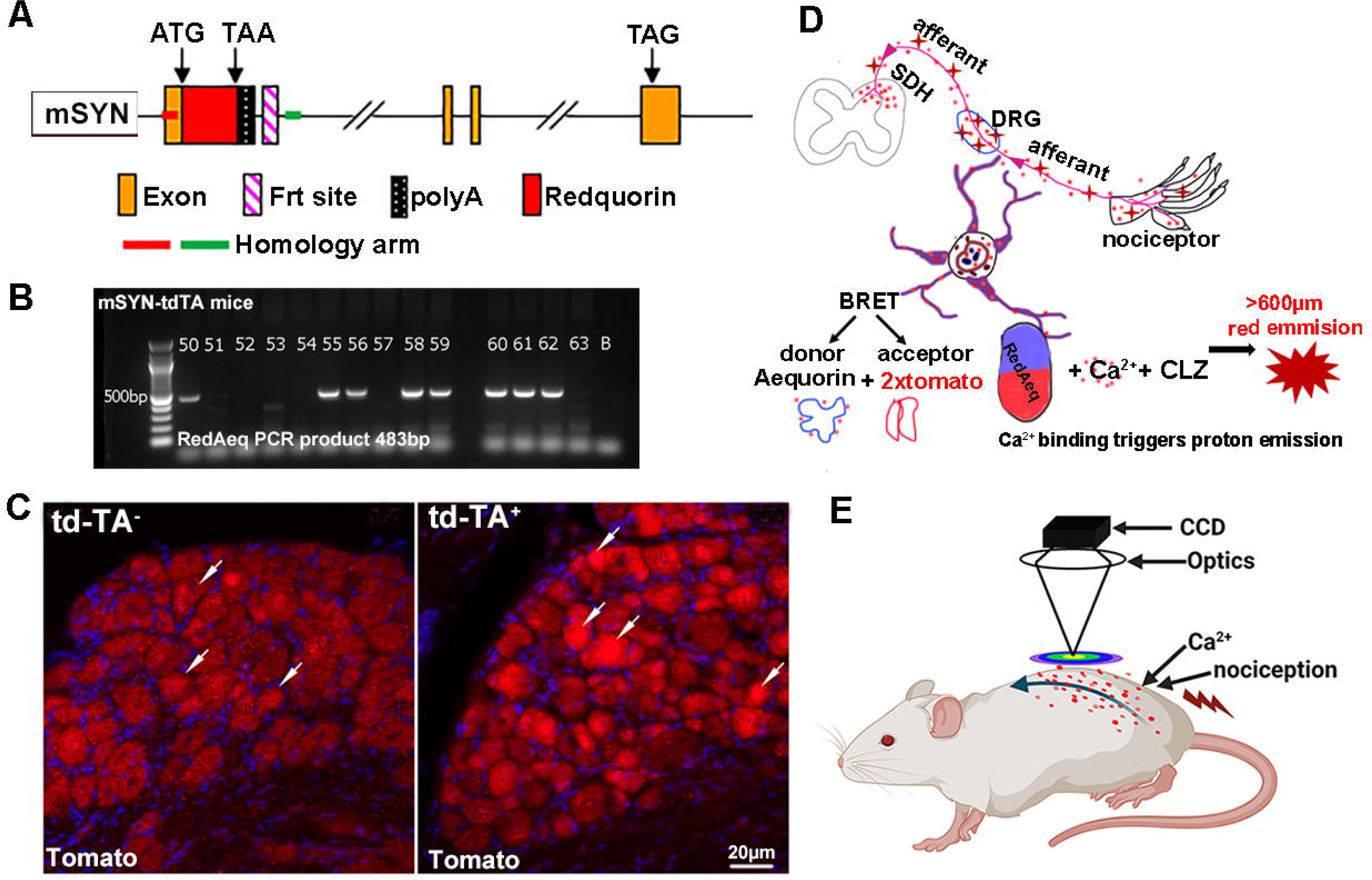
Design of nociceptive neuronal activity imaging (NNAI) mice. BAC, bacterial artificial chromosomes; BLI, bioluminescence imaging; BRET, bioluminescence resonance energy transfer; CCD, charge-coupled device; CLZ, coelenterazine; DRG, dorsal root ganglion; NNAI, Nociceptive Neuronal Activity Imaging; mSyn, mouse synapsin 1 promoter; Redquorin, tdTA, 2×tdtomato-aequorin. A. BAC mSyn -Redquorin construct. Redquorin, consisting of tdTA as reported^[2]^, is placed under the control of the mSyn1 in a BAC clone by homologous recombination. The BAC construct was used to generate the NNAI transgenic mouse. B. tdTA mouse pups were genotyped by PCR and those with the correct transgene product band 483bp, either homozygous or heterozygous were used for experimentation. C. The expression of the Redquorin transgene. DRG neurons of mSyn-tdTA^+^ mouse expressed td-tomato. Td-tomato immunopositive DRG neurons of mSyn-tdTA^+^ mouse (right), indicated by arrows, compared with arrows in litter mate tdTA^-^ mouse lumbar DRGs (left). D. Principle of NNAI mouse to produce BLI. The aequorin naturally occurring with blue light emission was modified by fusion with red emission tandem dimer (td) Tomato forms a chimera protein, which was put under the control of mSyn 1 promoter to restrict its expression in neurons. Coelenterazine (CLZ-*h*) is a nature substrate of aequorin with a role of oxygenizing aequorin; it was delivered by intrathecal injection (i.t.) prior to imaging. Nociceptive signal processing in sensory neurons elicited Ca^2+^ elevation. The Ca^2+^ binding to aequorin along with the presenting of CLZ-*h* triggers an oxidative decarboxylation of CLZ-*h* and the action results in photoprotein transformed from an excited aequorin (donor) to a 2xtd domato (acceptor) based on the principle of resonance energy transfer (BRET), which allows for energy transferring from the donor aequorin to the acceptor, the tdTomato, that emits bioluminescent photons. E. NNAI imaging with IVIS imager. The Redquorin released a long wavelength (≥600 µm) photon which is a red bioluminescent signal emitted from deep embedded neuron and nerve in NNAI mice and is imaged by IVIS imager® with a charge-coupled device camera aimed at mouse back.

**Fig 2.**
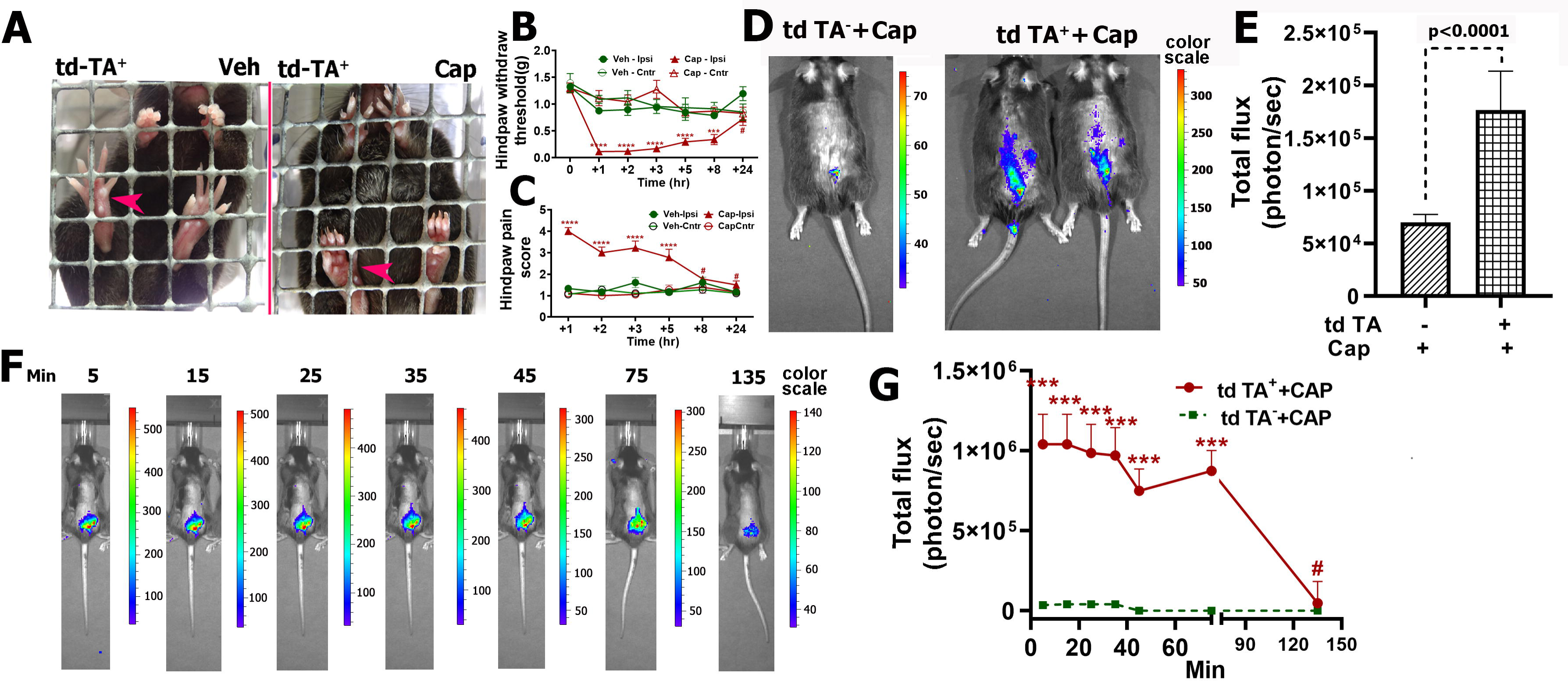
Measurement of capsaicin-induced nociceptive activity in pain circuits using NNAI mice. BLI, bioluminescent imaging; i.d., intradermal; NNAI, Neuronal Activity Imaging. A. Capsaicin-induced mechanical allodynia in the NNAI mouse. Mice were injected (i.d.) with capsaicin vehicle (100% alcohol: saline: Tween 80 2:1:7) or capsaicin 0.25% 5µl (A) in the ipsilateral hindpaw (red arrow). Two hours post capsaicin administration, mice manifested evident pain-elicited hindpaw postures (curled-up hindpaw), while control mice with vehicle injection exhibited a relaxed hindpaw posture. B. Capsaicin decreased mechanical allodynia expressed as hindpaw withdraw threshold (PWT) measured by von Frey test. Ipsilateral hindpaw showed a decreased PTW, compared with the contralateral in the same mice and ipsilateral hindpaw in the vehicle group. (paired student t test, p<0.0001, n =7 mice in both groups), with sex ratio 1:1. C. Quantification of hindpaw pain score after capsaicin i.d. Mice hindpaw postures were scored as described (paired student t test, p<0.0001, n=7 mice in each group). D. BLI images generated by neuronal activity-elicit Ca^2+^ rising. Photographic and luminescent images captured by the IVIS charge-coupled device (CCD) camera were overlaid. Capsaicin i.d. hindpaw showed significantly increased bioluminescent signals in the lumbar spinal regions (right) compared with the vehicle group(left), Images were acquired at 40 min after capsaicin injection. E. Quantification of BLI images. Bioluminescent photo density was quantified based on the total influx of photons using Cataloging & Browsing Tools. Parametric student *t* test, p<0.0001, n=8/group. F. Time course of bioluminescent signals induced by capsaicin. BLI images acquired at different time points after capsaicin injection. G. Quantitative temporal profiles of bioluminescent signals (n=6).

### Acquisition of ex vivo bioluminescent imaging (BLI)

The whole pain visualization imaging processes included tdTA mice genotyping, pain mouse model generation, and tdTA chimeric protein reconstitution with coelenterazine (CLZ-*h*). tdTA pain mice were prepared for *ex vivo imaging* and imaged with an IVIS Spectrum Imager (PerkinElmer, Shelton, CT) set at the “open” filter with the optical exposure time. The bioluminescent substrate XenoLight RediJect CLZ-*h* (Perkin Elmer part No. 760506) was used to reconstitute tdTA by intrathecal (i.t.) injection, as we previous described^[41; 55]^. Briefly, mice were initially anaesthetized with 3% isoflurane oxygen with a flow rate of 1.5L/min and maintained with a reduced isoflurane concentration at 2.5%. A 30.5-gauge stainless steel needle attached to a 25μL luer tip syringe (Hamilton, Reno, NV) was used for i.t. injection. The needle tip was placed in the gap between L5 and L6 at a ≤45° angle; a confirmation of successful intrathecal puncture was a tail twitch. The C2-modified native coelenterazine is hydroxyphenyl and aromatic. Using H to replace the original phenolic OH produces coelenterazine-*h*, which imparts a luminescent intensity with its aequorin complex reported to be 10–20 times higher than that of native coelenterazine, making this derivative a useful tool for measuring small changes in Ca^2+^ concentrations.

Formula: _26_H_21_N_3_O_2_. MW = 407.5.

The concentration of the ready-to-use CLZ-*h* is 150 μg/mL in propylene glycol/citrate. Our mouse optimized maximum dose was 0.15 μg (1 µl)/g body weight (gbw) with the route of i.t. injection. We used less than the recommended dose of CLZ-*h*30 μg /20gbw in mouse, as the CLZ-*h* was highly accurate in being delivered to the neuronal pain circuit. During and after CLZ-*h* injections, the anesthetized mouse was kept on a heated pad until recovery from anesthesia. Mice were imaged within 1hr post i.t. injection. During the imaging capture process, anesthetized mice were kept prone on a heated sample stage of the IVIS imager to maintain normothermia, and anesthetization was maintained by a mouthpiece connected to 2.5% isoflurane in oxygen with a flow rate of 1.5 L/min.

The IVIS® Spectrum Imager was used for image capture. The charge-coupled device (CCD) camera, with a raw signal range of 0 to 65,535, has the capacity to obtain a signal level of 600 to 61,000 counts, analog to digital counts (2^16^) ^[46]^. Images were captured by using the IVIS Acquisition Control Panel in luminescent mode that measures the light signal using an open filter in unit of photons with 5 min exposure time. A calcium-triggered bioluminescent image was captured in bioluminescence mode. Serial images were captured continuously within a 10 min interval. Standard Images were composed of two images: photographic + luminescent = overlay. The bioluminescent photon counts from the mouse back was captured in the region of interest (ROI) and was quantified by using fixed-size and fixed-position ROIs throughout the experiments. All images were measured by ROI measurements that come with the IVIS® Spectrum Imaging Series. Calibrated units were photons per second, representing the total flux radiating omni-directionally from a user-defined region. Measurement results were exported to an Excel spreadsheet.

### Mouse pain models

#### Capsaicin

tdTA mice were treated with two representative algogens, capsaicin and glycoprotein gp120 (gp120), to create two pain mouse models with distinct nociceptive mechanisms^[33; 51; 56]^. First, a capsaicin-induced acute pain model was generated as we have published^[41]^. Briefly, mice were anesthetized with isoflurane (2.5% for induction and 2.0% for maintenance) in an oxygen flow rate of 1.5L/min and placed in a prone position. For each mouse, 5μl of 0.25% capsaicin, dissolved in a vehicle of saline containing 20% alcohol and 7% Tween 80, was injected intradermally (i.d.) into the plantar region of the left hindpaw using a 30.5-gauge needle attached to a 10 μl Hamilton air-tight luer syringe. Mice injected with vehicle were used as controls. Simultaneously, mice injected i.t. with coelenterazine CLZ-*h* were submitted for imaging within 60 mins post injection.

#### gp120

A gp120 neuroinflammatory pain mouse model was generated as we described before^[56]^. The recombinant HIV-1 gp120Bal, Cat # ARP-4961 produced in HEK293 cells was provided by the NIH AIDS Research and Reference Reagent Program and stored at −80°C. At the time of injection, stock gp120 protein solution (1μg/lμL) was diluted with 0.1% bovine serum albumin in 0.1 M phosphate-buffered saline at a final concentration of 20ng/lμL. The amount of gp120 administered in mice was 5 μL/20gbw per mouse.

#### Neuropathic pain

A neuropathic pain tdTA mouse model was created by L5 spinal nerve ligation (SNL) conducted in Dr. Jin Mo Chung’s lab^[22]^. Female 4-5 month old tdTA mice were placed under anesthesia with isoflurane mixed with oxygen (3% induction and 2.5% maintenance). The lower back fur was shaved and a midline incision was cut. The muscles on top of the L5/6 vertebrae and the L6 transverse process were removed to get a good view of left L5 spinal nerve. The L5 nerve was ligated in its original location. The incision was sutured, and the anesthesia was stopped. Mice were returned to their home cage to recover. The same procedure was applied to the sham mice until a good view of the L5 spinal nerve was obtained, but without ligating it^[18; 53]^.

### Pain measurement

#### Hindpaw mechanical allodynia

Hindpaw mechanical allodynia was measured by means of the Von Frey test and the data expressed as hindpaw withdrawal threshold (PWT), as described before^[54]^. Briefly, prior to data collection, mice were habituated by restricting them for three days within a plexiglass box standing on a mesh floor continuously. On the test day, a calibrated von Frey filament (Stoelting, Wood Dale, IL) was applied perpendicularly to the central plantar area of the hindpaw, so that when the hindpaw was placed on the metal mesh floor, the filament started to bend like a hook and maintained contact for about one second. Positive nocifensive behaviors included vertically withdrawing or horizontally removing the hindpaw. The stimulation force was determined according to the Dixon up down method. The interval between each stimulation was ≥ 30 sec to minimize the effects of the previous stimulation. Stimulation was performed only when the mouse was in a resting and alert state, avoiding mice in the state of deeply sleeping, actively exploring, or grooming^[8]^. The experimenter was blinded to the group and treatment. Baseline was an average of three continuous days’ test results. The PWT value was calculated using an application coded according to the Dixon up down method and plotted with Prism 10.2.2 (GraphPad Software, La Jolla, CA). The PTW value was expressed as mean ± standard error.

#### Hindpaw relaxation measurement

We used the score of hindpaw relaxation as a indicator of spontaneous pain, as we described previously, based on the principle that spantaneous pain is associated with a decrease in hindpaw relaxation^[42; 54]^. Briefly, mice were habituated to the room prior. Then Von Frey testing was conducted, with spantaneous pain-related hindpaw curled-up postures scored and photographed for evaluation and documentation. Each hindpaw photo clearly included two hindpaws and five photos were taked at intervals of ≥15 secs. The pictured hindpaw postures were scored by a experimenter blinded to the experiemental group design, who counted the number of curled-up toes in both hindpaws. Scoring was: 0, no curled-up toes and the hindpaw flattly placed on the mesh floor, indicating no pain in the hindpaws; 1, one toe slightly curled up or bent; 2, two toes gently curled up or bent; hence, 5 was all toes tightly curled up and bent. The toe relaxation score of each hindpaw ranged from 0 to 5. Zero was the lowest score and 5 was the highest score. Each hindpaw score was an average of the ≥ 3 individual scores. The naïve or vehicle-only treated mice usually scored < 2.5. A higher score indicated more severe spontaneous pain.

## Results

### Development of tdTA mice with Redquorin gene construct

We designed an optical imaging strategy to measure spontaneous pain by monitoring nociceptive neuronal activity in pain circuits in mouse models. To this end, we sought to generate a transgenic mouse, for the purpose of NNAI, that would enable optical imaging of activity occurring in pain-processing neurons. Aequorin is a bioluminescence Ca^2+^-sensitive photoprotein which has been used to monitor Ca^2+^ elevation in various biological processes^[24]^. However, because the bioluminescence emitted from aequorin has poor tissue penetration due to a blue light wavelength of 450-470 nm, it is not suitable for imaging the pain circuits deeply embedded in tissues. Hence, a tdTA transgenic mouse was constructed by using a Redquorin gene construct^[2; 4]^. Redquorin is a hybrid fusion protein consisting of an aequorin and 2xtdTomato, so that Ca^2+^-induced blue light photon emitting from aequorin (the donor) are shifted to a red-light emission (>600 nm) in the acceptor tdTomato (>600 nm), based on the mechanism of BRET (**Fig. 1A**). Thus, the tdTA mouse has an enhanced ability of tissue photon penetration, as tissue is quite transparent to red light.

The tdTA mouse bearing the Redquorin construct was a BAC transgenic mouse. The Redquorin was placed under the control of the mSyn1 promoter via homologous recombination (**Fig. 1A**). mSyn1 is a neuronal-specific protein widely expressed in both the central and peripheral nervous system and concentrated in the synaptic region^[12; 29; 40]^. Thus, Redquorin expression and bioluminescence emission were restricted to the neuron This promoter enables long-term and stable expression of Redquorin exclusively to neurons which are functionally distinct, including small diameter sensory neurons with unmyelinated or thin myelinated (C- or 6) fiber or polymodal sensory neuron, all with a high threshold for pain transduction. Various nociceptive signal transmission neurons reside in the SDH^[14; 25; 52]^. The tdTA mice were generated by pronuclear BAC microinjection, the Redquorin gene was genotyped by PCR (**Fig. 1B**), and its expression was confirmed in DRG neurons by fluorescent immunostaining (**Fig. 1C**).

We used this BAC transgenic mouse (hereafter the tdTA mouse) in subsequent experiments to evaluate its potential usefulness in measuring spontaneous pain by IVIS imaging (**Fig. 1D**). We first used the tdTA mice to generate pain models, then performed the IVIS imaging. The collected IVIS images encompassed the mouse back and were well exposed to the IVIS CCD camera lens (**Fig. 1**, including the spinal regions; in particular, **1E**). The imaging visualized the lumbar spinal region, which is most responsive to ipsilateral hindpaw noxious stimulation, and was well exposed to the IVIS CCD camera lens (**Fig. 1D**). This imaging approach can objectively capture the neuronal activity-elicited longwave bioluminescent photon emitted from Redquorin borne by tdTA mice (**Fig. 1D, E**).

### Imaging capsaicin-induced spontaneous pain in tdTA mice

To evaluate the efficacy of tdTA mice in generating BLI in an acute pain model, we utilized an intradermal injection of capsaicin (0.25% 5µl) in the hindpaw of tdTA mice. The capsaicin-injected mice developed mechanical allodynia as measured by the Von Frey test immediately post-injection, and the lowest mechanical threshold peaked at 1-3 hr thereafter (**Fig. 1B**). Concurrently, the hindpaws exhibited decreased relaxation postures (**Fig. 1A, C**)^[42; 55]^. Observations confirmed that the mice developed both evoked and spontaneous pain following i.d. capsaicin administration.

To prepare mice for BLI imaging, in addition to capsaicin injection, CLZ-h was also i.t. administered simultaneously. At 1 hr post-CLZ-h i.t., BLI images were acquired using an IVIS imager. The acquired IVIS images revealed a remarkable increase in photon emission in response to capsaicin in tdTA^+^ mice (**Fig. 1D, E**), whereas the wild type (WT) and littermates (tdTA^-^) displayed little bioluminescent signal (**Fig. 1D, E**). Photon total flux was significantly higher in tdTA^+^ mice compared to their tdTA^-^ counterparts (**Fig. 1E**).

Subsequently, we investigated the temporal profiles of BLI following capsaicin administration and Redquorin reconstitution. Serial BLI was captured at certain time intervals post-capsaicin injection (**Fig. 1F**). Quantitative analysis revealed stable emission with minor decay up to 75 min post-capsaicin administration, followed by a pronounced decline at 135 min (**Fig. 1F, G**). Because our behavioral testing showed that mice still manifested spontaneous pain at its peak at 120 min post-capsaicin injection (**Fig. 1A, B**), the rapid decay in photon emission around this time was due to the decrease in CLZ-*h* in the spinal cord over time, rather than a decrease in intensity of spontaneous pain. This distinct temporal window for BLI imaging, maintaining efficacy for up to 75 min, is pivotal for optimizing the experimental design using the tdTA mice.

We developed and optimized a BLI imaging protocol that included two separate injections. The first injection was used to create pain models, and the secondary injection was used to reconstitute the Redquorin with CLZ-*h*. After successful pain induction followed with aequorin reconstitution, the detectable BLI can be maintained up to 75 min post CLZ-*h* administration without obvious decay. This setting provided the maximal time window to take the BLI and the duration allows full nociception development that fits with the capsaicin acute pain model. We next adapted this imaging protocol to evaluate sub-acute (gp120) and chronic pain (SNL), and to assess the analgesic effect of gabapentin analgesia in SNL mice. The advantage of the BLI process is that it is virtually non-invasive and automated, avoiding any experimenter-animal interaction that may affect the nociceptive test experimental readout.

### Imaging HIV-associated pain in tdTA mice

To test the tdTA mouse’s capability to discern inflammatory pain, we utilized an HIV-associated pain model as we previously described^[55; 56]^. This model was generated by i.t. injection of gp120, an HIV-1 envelope protein implicated in the development of HIV-associated pain^[41; 42; 56]^. tdTA^+^ mice that received a single i.t. dose of gp120 (100ng/20gbw) subsequently developed mechanical allodynia at 18 hr post gp120 i.t. This allodynia was maintained for up to 4 days (**Fig. 3A, B**).

**Fig 3.**
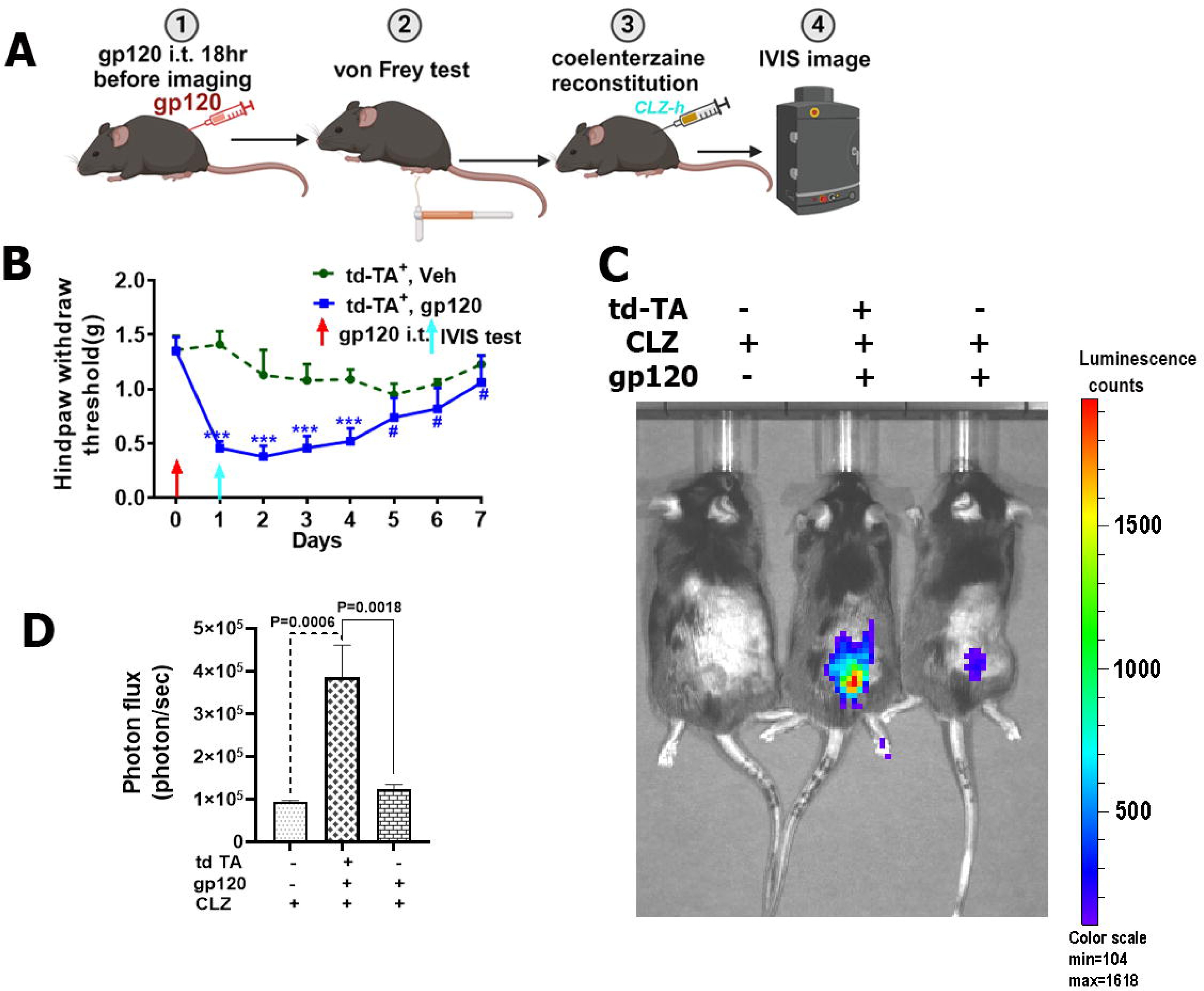
Measurement of HIV-1 gp120-induced nociceptive neuronal activity in pain circuits using NNAI mice. CLZ-*h,* tdTA chimeric protein reconstitution with coelenterazine; gbw, g body weight; i.t., intrathecal; IVIS, in vivo imaging system; NNAI, Neuronal Activity Imaging. A. A scheme of experimental processes. ① gp120 administration, i.t. gp120 100ng/20gbw. ② At 18 hr post gp120 i.t., Von Frey test was conducted. ③ Photoprotein substate CLZ-*h* was administrated via i.t. injection at a dose of 1µl/1gbw. ④ IVIS imaging. B. gp120 induced mechanical allodynia in NNAI transgenic mice. Adult male NAI transgenic mice received a single i.t. of gp120 (i.t.,100ng/20gbw), and von Frey tests were performed at 18 hr post i.t. injection and daily thereafter. The vehicle group was i.t. with equal amount of heat-inactivated gp120 and used as vehicle control. *Two-way* ANOVA tests with Tukey’s multiple comparisons post hoc tests were performed (***, P<0.001, n=6), with sex ratio 1:1. C. IVIS imaging at the lumbar spinal level of NNAI transgenic mice after gp120 injection (i.t.). Adult (3-4 month) tdTA^+^ and tdTA^-^ littermates were used in the experiment. D. Quantification of IVIS bioluminescent images. One-way ANOVA with Tukey’s multiple comparison tests were used to compute *p* values (n=6).

For nociceptive BLI imaging, we allowed mice to fully develop pain, reaching peak PWT at 18 hr; at that time, BLI imaging was taken (**Fig. 3A**). tdTA^-^ mice administered CLZ-*h* (i.t.) only, or gp120 with CLZ-*h* (i.t.) together, exhibited little signal in the lumbar area. In contrast, tdTA^+^ mice administered gp120 and CLZ-h together (i.t.) showed a significantly elevated bioluminescent photon flux (3.86×10^5^, 3.8-fold increase, n=6, p=0.0025, one-way ANOVA with Tukey’s multiple comparisons test) (**Fig. 3C, D**). These results suggest that tdTA^+^ mice can visualize inflammatory pain.

### Imaging neuropathic pain in tdTA mice

Next, we sought to test the potential use of tdTA mice in imaging neuropathic pain in SNL mice. To this end, we generated a SNL neuropathic pain model using tdTA mice, with tdTA^-^ littermates as controls^[22]^. These SNL mice developed evoked mechanical pain in the ipsilateral hindpaw, as shown by von Frey tests (**Fig. 4A**), and spontaneous pain, as measured by scoring hindpaw relaxation gestures (**Fig. 4B, C**).

**Fig 4.**
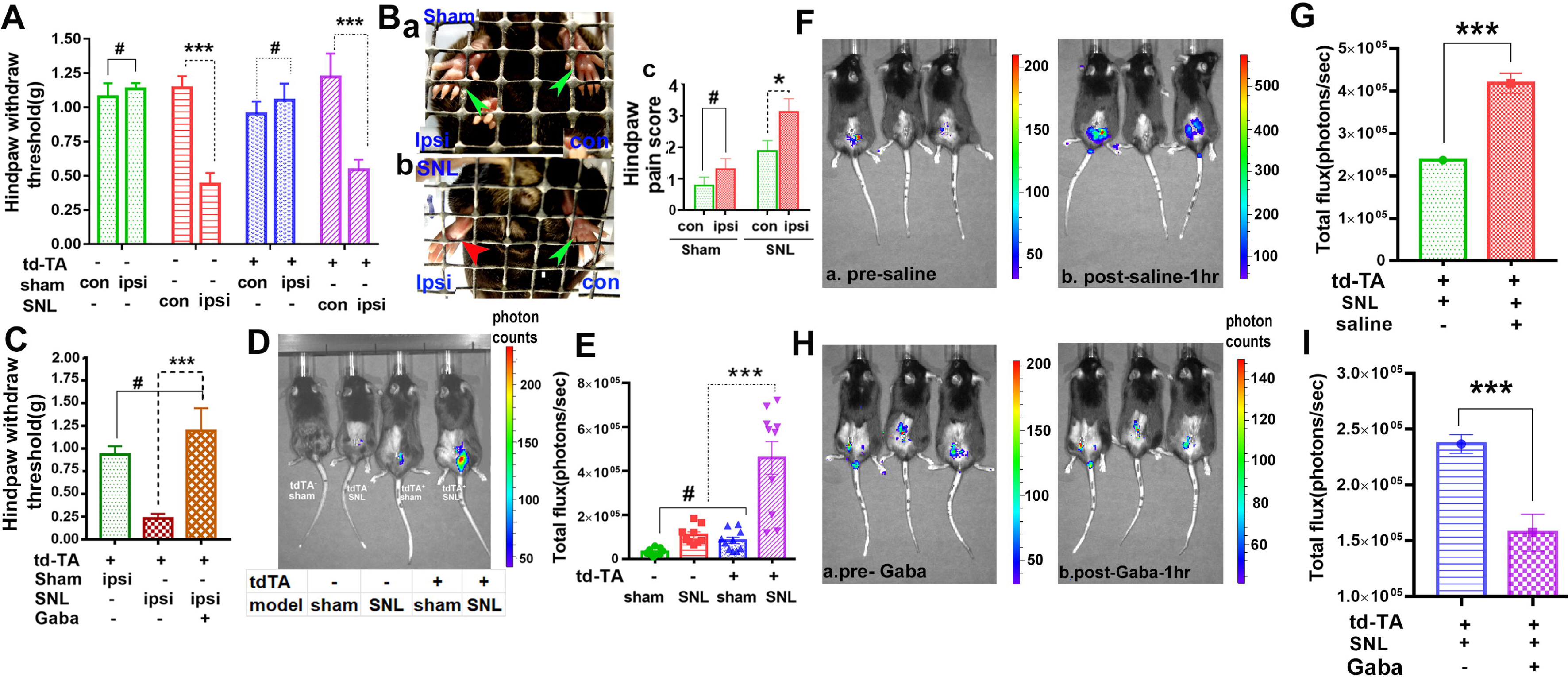
Measurement of nociceptive neuronal activity in pain circuits in a neuropathic pain NNAI mice. BLI, bioluminescence imaging; i.p., intraperitoneal; NNAI, Neuronal Activity Imaging; PWT, hindpaw withdrawal threshold; SNL, sciatic nerve ligation. A. Development of mechanical allodynia after SNL in NNAI transgenic mice. von Frey tests showed that tdTA^+^ (purple) and tdTA^-^ (red) mice manifested a decreased PWT in the ipsilateral hindpaw at 6 days post SNL surgery, whereas both tdTA^-^ (green) and tdTA^+^ (blue) mice with sham surgery, the contralateral and ipsilateral hindpaws showed no significant difference in PWT. Sham tdTA^-^ mice, n=6; SNL tdTA^-^ n=6; Sham tdTA^+^ n=6; SNL tdTA^+^ n=7, with sex ratio 1:1, *** p<0.001, two-way ANOVA with Tukey test for multiple comparisons. B. SNL induced mechanical allodynia in the ipsilateral hindpaw of the NNAI (tdTA^+^) mouse at 6 days post SNL surgery. **a**. Both contralateral and ipsilateral hindpaws (two green arrows) of sham mice exhibited a relaxed posture, indicating no spontaneous pain in the hindpaws. **b**. The ipsilateral hindpaw of SNL mice manifested evident spontaneous pain-related curled postures (red arrow), while the contralateral hindpaw of SNL mice showed a relaxed gesture (green arrow)^[55]^. **c**. Quantification of hindpaw pain scores. #, no significance, sham group, n=6; SNL group, n=7, with sex ratio 1:1. * p<0.05, two-way ANOVA with Tukey test for multiple comparisons. C. The analgesic effect of gabapentin on SNL-induced mechanical allodynia in NNAI mice. One-time i.p. gabapentin reversed the decreased PWT at 9 days post SNL surgery; one hour post gabapentin administration (100mg/kg i.p), the SNL allodynia in the ipsilateral hindpaw was blocked, as shown by a significantly elevated PWT compared with SNL mice treated with saline (i.p). One-way ANOVA test, ***<0.001, sham, n=6; SNL, n=7; SNL mice treated with gabapentin, n=8, mice with sex ration 1:1. D. BLI imaging of SNL NNAI mice at day 7^th^ post-surgery. All sham and SNL tdTA^-^ and tdTA^+^ mice were i.t. injected with 1 μl/gbw CLZ-*h* and imaged 40 min post-injection in the IVIS imaging chamber. The injection and imaging were performed under anesthesia by inhaling 2% isoflurane in oxygen. Three images were taken continuously at intervals of 10 min and the total influx was the average of the three images. The representative tdTA^-^ sham and SNL mice presented only very low background signals. The representative tdTA^+^ sham mouse showed more background signals compared with tdTA^-^ sham and SNL mouse. Only the tdTA^+^ SNL mouse displayed a significantly increased bioluminescent signal. E. Quantification of D. The total flux of photons from tdTA^-^ sham, tdTA^-^ SNL, and tdTA^+^ sham mice all showed low total flux and no significant differences between the three groups (# p>0.05, n=9, each group, sex ration 1:1). When the three groups compared with tdTA^+^ SNL mice, the latter (n=9, sex ration 1:1) showed significantly higher bioluminescence flux, compared with the other three groups (***p<0.001). One-way ANOVA test was used to evaluate significance among means and Tukey’s multiple comparison test used to calculate the means between groups F & G. Saline treatment showed no analgesic effects on tdTA^+^ SNL mice by comparing the BLI pre- and post-saline i.p. injection. **a.** Representative BLI image of tdTA^+^ SNL mice prior to saline treatment. **b.** Saline (i.p.) treated mice showed a significantly increased bioluminescent signal at 1 hr post-saline, n=7, mix sex, student t test, ***p<0.001. H & I. Analgesic effects of gabapentin on bioluminescent signals in tdTA^+^ SNL mice. **Ha.** A representative BLI image of tdTA^+^ SNL mice prior to gabapentin treatment (pre-Gaba). **Hb.** tdTA^+^SNL mice were treated with gabapentin (i.p. 100mg/kg in 0.2 ml saline). At 1 hr post- treatment, the BLI image showed a significantly decreased bioluminescent signal compared to the same group (pre-Gaba) of tdTA^+^ SNL mice. **I**. At 1 hr post-gabapentin injection, tdTA^+^ SNL mice showed a significantly decreased bioluminescent signal. n=7, mix sex 1:1, student t test, ***p<0.001.

We then captured IVIS imaging of the tdTA^+^SNL and tdTA^-^ mice at 7 days post-surgery. The BLI showed significantly higher photon flux emission only from tdTA^+^SNL mice compared to three groups of control mice: tdTA^-^ sham, tdTA^-^ SNL, and tdTA^+^ sham (p<0.0001) (**Fig. 4D, E**). The results demonstrated that Redquorin-mediated photon emission acquired by the IVIS imager is sufficient to detect and visualize the pain circuitry activation caused by nerve injury from SNL.

To further investigate whether photon emission can be used to evaluate the analgesic effect in drug development, we tested the effect of gabapentin, an analgesic widely used to manage neuropathic pain in clinics and in animal models^[5; 47; 50]^. We observed that gabapentin administration (100mg/kg, intraperitoneally) effectively reversed mechanical hypersensitivity in tdTA^+^ SNL mice at 1 hr post-gabapentin injection (**Fig. 4C**). Consistent with these results, we observed in BLI that, compared with saline-treated mice (**Fig. 4F & G**), gabapentin administration significantly inhibited the increase in photon emission in tdTA^+^ SNL mice (**Fig. 4H & I**) (p<0.0001, *t*-test). These data suggest that the analgesic effect of gabapentin on neuropathic pain can be measured and visualized by BLI in tdTA^+^ mice.

### Visualization efficiency of NNAI mice

We observed different negative and successful rates with different pain models (**Table 1**). The positive rate of BLI of capsaicin-treated tdTA^+^ mice was 53.19% vs a negative rate of 46.81%; in C57BL6 mice (WT, do not carry tdTA mutant gene) the negative rate was 92.86% and in tdTA^-^ mice was 100%. The high negative rate in capsaicin-treated WT and tdTA^-^ mice makes the NNAI mouse a reliable tool to detect capsaicin-induced pain with the mechanism of nociceptor activation triggered by rising neuronal calcium levels.

**Table 1.** Successful rates of BLI with different pain models. BLI, bioluminescent imaging; SNL: spinal nerve ligation; tdTA: tandem-dimer tomato-aequorin; wt, wild type.

| Capsaicin | total mice tested | mice (-) | % | mice (±) | % | mice (+) | % |
| --- | --- | --- | --- | --- | --- | --- | --- |
| Wt | 14 | 13 | 92.86% | 1 | 7.14% | 0 | 0.00% |
| tdTA<sup>-</sup> | 8 | 8 | 100.00% | 0 | 0.00% | 0 | 0.00% |
| tdTA<sup>+</sup> | 47 | 22 | 46.81% | 0 | 0.00% | 25 | 53.19% |

| gp120 | total mice tested | mice (-) | % | mice (±) | % | mice (+) | % |
| --- | --- | --- | --- | --- | --- | --- | --- |
| tdTA<sup>-</sup> | 9 | 5 | 55.56% | 0 | 0.00% | 4 | 44.44% |
| tdTA<sup>+</sup> | 21 | 7 | 33.33% | 0 | 0.00% | 14 | 66.67% |

| sham/SNL | total mice tested | Sham |  |  |  | SNL |  |  |  |  |
| --- | --- | --- | --- | --- | --- | --- | --- | --- | --- | --- |
|  |  | + | % | - | % | total mice tested | + | % | - | % |
| tdTA<sup>-</sup> | 6 | 0 | 0.00% | 6 | 100.00% | 8 | 0 | 0 | 8 | 100.00% |
| tdTA<sup>+</sup> | 10 | 0 | 0.00% | 10 | 100.00% | 34 | 32 | 94.12 | 2 | 5.88% |

The HIV-1 gp120 sub-acute tdTA^+^ pain mice model showed a 66.67% positive rate of BLI compared with a 44.44% positive rate in tdTA^-^ mice. Even though the photon signal exhibited a very low photon influx, as indicated in **Fig. 3**, we still count it as positive. The gp120-treated tdTA^-^ mice that showed a 44.44% low photon flux we counted as positive. This is a known phenomenon in BLI that genotyped negative mice show a positive background instead of being completely negative^[19]^. The 53.19% positive rate in capsaicin-treated and 66.66% positive rate in tdTA^+^ mice indicate successful BRET efficiency and BLI imaging.

Fully 66.67% of gp120 tdTA^+^ mice showed a positive BLI, suggesting that tdTA mice can successfully show gp120-induced inflammatory pain. We tested the mice at 18 hr post gp120 i.t., when gp120-induced mechanic allodynia reached the lowest hindpaw withdraw threshold. The gp120 dose we used approximated level found in the postmortem SDH of HIV-infected patients with chronic pain (HIV+ PAIN+)^[56]^. The BLI was taken at the time when mice showed the lowest mechanical threshold and peak time of allodynia. Pain via gp120 is due to gp120 binding to its CD44 receptor and co-receptor CXCR4 and CCR5 in microglia or astrocytes^[26]^, with the later secretory chemokines and cytokines initiating neuroinflammation. The gp120-elicited BLI suggests that the NNAI mice can detect neuroinflammation induced calcium rising in neuronal.

## Discussion

Currently-used popular methodologies, the Von Frey, Hargreaves (heat) and cold tests, are predominantly designed to measure nociception responses to noxious stimulations^[13]^. Although useful for measuring different modalities of pain, such tests cannot measure spontaneous pain, which is commonly experienced in human patients and in animals. Various behavioral paradigms have been developed in recent years to address this unmet need, including conditioned preference place (CPP)^[56]^, weight bearing foot printing[13], flinch and licking^[27]^ gait change, and paw gestures^[42; 55]^. However, these approaches, although useful, are often confounded by learning (e.g., CPP) and subjective interpretation (e.g., flinch and licking, paw gestures, etc.). In this study, we developed a BAC transgenic mouse, the NNAI generating tdTA mouse, by expressing Redquorin^[2]^ under the control of the synapsin 1 promoter. Using the property of Redquorin to emit longwave red light in response to Ca^2+^ elevations in active neurons, we can measure neuronal activity in the pain circuits as a nociceptive biomarker.

In the tdTA SNL neuropathic pain mouse model, tdTA^-^ and tdTA^+^ sham mice, as well as tdTA^-^ SNL mice, show 100% negative BLI, while only the tdTA^+^ SNL mice showed a high positive rate of BLI (94.12%), with only 5.88% of mice showed an acceptable negative rate. This result makes NNAI mice an extremely promising approach to detect SNL-induced neuropathic pain.

To target the spinal cord and DRG pain circuit, we optimized the CLZ-*h* dose and administration route according to the instructions of the ready-to-use coelenterazine (PerKinElmer). Our results suggest that the CLZ dosage and administration route must be optimized based on which organ or tissue the BLI is designed to target. Another reason for a negative BLI is that the Redquorin photoprotein expression may be silenced following multiple years of breeding. This can be avoided by cryopreservation and re-derivation of an earlier generation of tdTA^+^ mice sperm and embryo, when necessary.

The effect of transgene silencing may be the underlying cause of negative BLI in tdTA^+^ mice. Transgene silencing has been observed in many mammalians cell-based transgene applications, as well as in transgene cargos carried by different viral tools and vectors. The capsaicin and gp120 tdTA^+^ pain mice models showed a 46.81% and 33.33% tdTA^+^ negative rate of BLI, respectively (Table 1). We double checked the genotyping of the negative mice in which silence occurred, despite its encoded

DNA remaining present with correct tdTA^+^ genotyping. Several mechanisms can cause transgene silencing, most commonly DNA methylation and histone modifications. The gene silencing usually can gradually decrease the relative levels of transgene expression, but loss of expression is never complete^[7]^. Therefore, accurately estimating when and what percentage of tdTA^+^ mice lose the capacity to generate BLI is difficult. One solution is to retrieve the tdTA^+^ mice from early generation cryopreserved mouse embryos with the right Redquorin expression, as proven by BLI imaging. Using an early generation NNAI mouse embryo to build clones and using it for BLI will improve the positive rate^[36]^. Two common mechanisms of transgene silencing and reactivation are: 1) promoter methylation, which occurs with the methylation of cytosine residues in CpG sequence by the cytosine DNA methaltrasferase enzyme, which can be reversed with methyltransferase inhibitor 5-aza-deoxycytidine^[44; 48]^; and 2) a post-transcriptional modification known as histone acetylation, which occurs when an acetyl group is added to a lysine residue on the tail of a histone by high level histone deacetylase, which can be prevented by a histone deacetylase inhibitor, sodium butyrate^[28]^ ^[11]^.

Our data indicates that the NNAI mouse offers a novel imaging approach to visualize non-evoked spontaneous pain in different pain mouse models and may provide a universal methodology to measure different types of pain associated with various disease conditions. In addition to its potential in measuring pain in living animals, this approach may be useful for large-scale testing of the efficacy of analgesic candidates in preclinical studies. It is also expected to find application in measuring neuronal activity in neural circuits involved in other neurological conditions such as epilepsy. Overall, our results suggest that the NNAI mouse provides a novel and innovative approach to detect spontaneous pain in pain models with different nociception mechanisms. For capsaicin and gp120 pain mouse models, BLI still needs further optimization to boost its positive rate and make the NNAI mouse a more promising approach in spontaneous pain detection.

In conclusion, we developed an imaging approach to objectively measure spontaneous pain in mice. Further optimizing of this system may lead to improvement of the NNAI mouse that has the potential for broad application in both basic and translational studies of pain and other neurological disorders.

## Acknowledgements

The authors have no conflicts of interest to declare. We gratefully acknowledge the assistance of Dr. Igor Patrikeev for all BLI imaging capture; Drs. Jun-Ho La’s and Jin Mo Chung’s lab for SNL surgery; and Dr. Sarah Toombs Smith for her reviews and editing the manuscript. This work was supported by pilot grants from the University of Texas Medical Branch Center for Interdisciplinary Research in Women’s Health, Sealy Center on Aging, and the Claude D. Pepper Older Americans Independence Center (P30 AG024832) and NINDS R01NS122571 (SY), as well as NIH grants R01NS079166, R01DA057195, R01DA050530 (SJT) and NINDS R01NS131507 (DPG).

## Abbreviations

BAC: bacterial artificial chromosomes
BLI: bioluminescent imaging
BRET: bioluminescence resonance energy transfer
CCD: charge-coupled device
CLZ-*h*: coelenterazine *h*
CPP: conditioned preference place
DRG: dorsal root ganglion
FRT: flippase recognition target
gbw: g body weight
gp120: glycoprotein gp120
i.d.: intradermal
i.t.: intrathecal
i.t.: intravenous
IVIS: in vivo imaging system
mSyn1: murine synapsin 1 promoter
NNAI: Nociceptive Neuronal Activity Imaging
OA: osteoarthritis
PWT: hindpaw withdrawal threshold
RA: rheumatoid arthritis
ROI: region of interest
SNL: spinal nerve ligation
SDH: spinal cord dorsal horn
tdTA mouse: mouse bearing 2x tandem dimer Tomato Aequorin
WT: wild type C57BL/6 mouse.

## Authors’ contributions

SY: Conceptualized and designed all the experiments, performed experiments, did data analysis and interoperation, and prepared the manuscript. JL: Mouse genotyping, performed pain behavior tests, prepared mouse for imaging, and performed data analysis. JP, mouse genotyping and pain behavior tests. JW: SNL mouse surgery. DG: Supervised genotyping and data interpretation. MM: Supervised the BLI imaging. SJT: tdTA mouse conceptualization and design.

Development of Nociceptive Neuronal Activity Imaging (NNAI) mice to visualize spontaneous pain, by transferring nociception-eliciting calcium rising to bioluminescence image.

